# Maturation of Sleep EEG Complexity in Preterm Newborns: Insights from Lempel-Ziv and Joint Lempel-Ziv Analyses

**DOI:** 10.64898/2026.08.10.742446

**Authors:** Andrea Devera, Magaly Catanzariti, Mariana Legnani, Carina Mezquita, Joaquín González, Luis Urban, Heber Hackembruch, Fernanda Blasina, Pablo Torterolo, Diego M. Mateos

## Abstract

The development of the sleep-wake cycle reflects the progressive structural and functional maturation of the brain. However, the organization of neural dynamics during prematurity remains incompletely understood. In this study, we analyzed the EEG from 54 polysomnographic recordings obtained from 39 preterm infants, grouped according to postmenstrual age (PMA) into three categories: 30-31, 32-33 and 34-35 weeks. Lempel-Ziv Complexity (LZC) and Joint Lempel-Ziv Complexity (JLZC) of the electroencephalogram (EEG) were analyzed during active sleep (AS), quiet sleep (QS), and indeterminate sleep (IS).

LZC computed from the raw, unfiltered recordings were significantly higher during QS than during AS and increased with PMA during AS. To further refine the analysis, LZC was also evaluated separately in the low-frequency (1-15.5 Hz) and high-frequency (16-30 Hz) EEG bands. In the low-frequency band, LZC was consistently higher during QS than during AS, an effect that was most pronounced in more immature groups. Furthermore, LZC increased with maturation particularly during AS. Sleep-state comparisons of LZC in the high-frequency EEG band also revealed higher values during QS than during AS across all PMA groups. Moreover, in contrast to the low-frequency band, LZC progressively decreased with advancing PMA both in AS and QS, suggesting that the neural mechanisms underlying low- and high-frequency EEG activity follow distinct maturational trajectories. Interestingly, larger LZC in the temporal cortex and interhemispheric differences were detected in the 32-33 PMA group. On the other hand, JLZC analysis revealed greater joint spatiotemporal dynamics across EEG channels during QS than during AS, with consistently higher JLZC values in temporal regions and lower in occipital regions.

Together, these findings show that these complexity metrics distinguishes sleep states and captures maturational changes in EEG activity in preterm infants. These results provide novel insights into early brain development and suggest potential quantitative biomarkers of neonatal brain maturation.

## 1. Introduction

Preterm birth is a major public health issue affecting more than 13 million babies worldwide each year, and preterm birth complications are the leading cause of death among children under 5 years old (World-Health-Organization, 2023). Although advances in neonatal care have markedly improved survival rates, a detailed understanding of the developing brain, both in healthy and clinically complicated preterm infants populations remains a priority.

Human brain development begins during the prenatal period and continues throughout the first two decades of life, with sleep playing a critical role in this maturational process (Riggins et al., 2024). During the third trimester of pregnancy, the brain undergoes accelerated developmental changes, including synaptogenesis, myelination, dendritic pruning, and the maturation of thalamocortical and corticocortical pathways (Kostovic et al., 2019;Kratimenos et al., 2025). Studies in preterm infants have consistently shown that sleep predominates during this stage of development (Parmelee et al., 1967;Curzi-Dascalova et al., 1988).

Sleep in newborns exhibits an immature organization, characterized by active sleep (AS), considered an immature form of rapid eye movement (REM) sleep; quiet sleep (QS), an immature form of non-REM (NREM) sleep; and indeterminate or undifferentiated sleep (IS), which shares characteristics of both AS and QS (Andre et al., 2010;Grigg-Damberger, 2016). The proportion of time spent in these sleep states changes as gestational age advances (Dereymaeker et al., 2017). According to our recent work (Devera et al., 2023;Devera et al., 2026) and “classical” studies (Curzi-Dascalova et al., 1988), AS predominates in preterm newborns, occupying most of the recording time; on average around 70% between 29 and 39 weeks of postmenstrual age (PMA), whereas QS accounts for approximately 18%. IS represents 8% of total sleep time and shows a marked tendency to decrease with increasing PMA. In contrast, wakefulness (W) occupies only a very small proportion of time (approximately 1%), although it progressively increases with maturation. It is important to highlight that AS not only occupies most of the time in preterm newborns, but also plays a critical role in brain maturation, as demonstrated in previous studies in both animals and humans (Curzi-Dascalova et al., 1993;Mirmiran, 1995;Mirmiran et al., 2003;Bertelle et al., 2005;Peirano and Algarin, 2007;Frank, 2017;Blumberg et al., 2022;de Groot et al., 2024). AS has also been proposed to play a critical role in the development of cognitive functions. In this regard, Hobson suggested that AS, and later REM sleep, may contribute to the emergence of a primordial form of consciousness, referred to as “protoconsciousness” (Hobson, 2009).

The electroencephalogram (EEG) during sleep also undergoes profound maturational changes throughout prematurity (Scher, 2008;Andre et al., 2010;Tsuchida et al., 2013;Wallois et al., 2021). While AS is characterized by continuous EEG activity, QS exhibits a discontinuous burst-quiescence pattern, with interburst intervals progressively decreasing as PMA advances (Andre et al., 2010;De Wel et al., 2021). Again, IS shares features of both AS and QS and is distinguished primarily by peripheral biosignals. However, detailed qualitative analyses of the EEG organization during sleep in preterm infants remain limited and inconclusive.

A detailed analysis of EEG activity in preterm infants is essential not only for understanding early brain maturation, but also for determining how continuous exposure to the unfavorable extrauterine environment of the neonatal intensive care unit (NICU) may influence this developmental process and subsequent long-term outcomes (Burlando et al., 2026). Preterm infants are exposed to multiple sensory stimuli, including noise, light, painful procedures, and caregiving interventions, all of which may disrupt sleep organization and brain electrical activity (Park, 2020).

Due to the complex nature of EEG signals, traditional neuroscience approaches that divide the spectrum into discrete frequency bands fail to fully capture the complexity of cortical electrical activity, especially during this developmental period. In contrast, the field of nonlinear dynamics has developed measures and models that account for the complexity of biological systems and their emerging interactions. The thalamocortical network dynamics, which are the basis of the EEG (Lopes da Silva, 2010), change during development (Wallois et al., 2021), as well as in the transition from conscious to unconscious states in adult individuals (Torterolo et al., 2019). These changes are captured by EEG-based measures of neural complexity (Schartner et al., 2015;De Wel et al., 2017;Mateos et al., 2017;Mateos et al., 2018;De Wel et al., 2019;Gonzalez et al., 2019;Moser et al., 2019;Sarasso et al., 2021;Gonzalez et al., 2022;Gonzalez et al., 2023a;Mondino et al., 2024).

In the present study, EEG complexity in preterm newborns was assessed using one of the most widely studied measures of neural complexity, Lempel-Ziv complexity (LZC) (Lempel and Ziv,1976). This metric, which quantifies the number of distinct substrings or patterns within a sequence offers several advantages for EEG analysis. It quantifies temporal pattern diversity directly from the signal, requires few assumptions about underlying dynamics and has been shown to be sensitive to changes in brain state and maturation in experimental and clinical settings.

To provide a more comprehensive characterization of preterm EEG activity, we also applied a multidimensional extension of LZC, namely joint Lempel-Ziv complexity (JLZC). Unlike conventional LZC, JLZC simultaneously quantifies the complexity of multiple signals, thereby capturing how their interactions contribute to the overall complexity of the whole system (Zozor et al., 2005). Therefore, the aim of the present study was to evaluate LZC and JLZC during AS, QS, and IS in EEG recordings from preterm newborns between 30 and 35 weeks of PMA.

## 2. Methodology

This study was conducted in the NICU of the Hospital de Clínicas, School of Medicine, Udelar, Montevideo, Uruguay, from November 2019 to March 2024. Ethical approval was obtained from the Clinical Research Ethics Committee of the same institution.

### 2.1. Population study

Eligible participants were preterm newborns with 30 to 35 weeks of PMA admitted to the NICU and clinically stable at the time of recording. Exclusion criteria were malformations of the central nervous system (CNS), intracranial hemorrhage, use of antiseizure medication or CNS-depressant drugs and chromosomal/congenital abnormalities. Parents of eligible preterm infants were invited to participate in the study. Written informed parental consent was obtained for all infant participants. Maternal clinical data related to pregnancy and delivery were obtained from the medical records. The subjects included in the present study represent a selected subset, based on PMA, from a larger cohort (Devera et al., 2026).

### 2.2. Polysomnographic recordings

Ninety-minute polysomnographic recordings, including EEG, chin (mylohyoid) and deltoid electromyography (EMG), electrooculography (EOG), electrocardiography, respiratory effort, and synchronized audio-video monitoring, were acquired using a Nihon Kohden EEG-9100 system. Data were acquired at a sampling rate of 500 Hz with 16-bit resolution and analog filtering between 0.5 and 70 Hz. Prior to electrode placement, the scalp was gently prepared with Nuprep® gel to reduce impedance. EEG activity was then recorded using disposable Ag/AgCl scalp electrodes (7-mm diameter) fixed with electroconductive paste. Electrodes were positioned according to the conventional reduced international 10-20 system, covering the frontal (F3, Fz, F4), central (C3, Cz, C4), temporal (T3, T4), and occipital (O1, O2) regions (Figure 1A, B), with additional electrodes placed over the mastoids (A1 and A2). The equipment acquired the signals referenced to C3-C4. The recordings were performed at 1:00 PM following feeding to facilitate spontaneous sleep onset in a temperature-controlled environment. In a subset of infants, recordings were repeated at three-week intervals during hospitalization until discharge.

**Figure 1.**
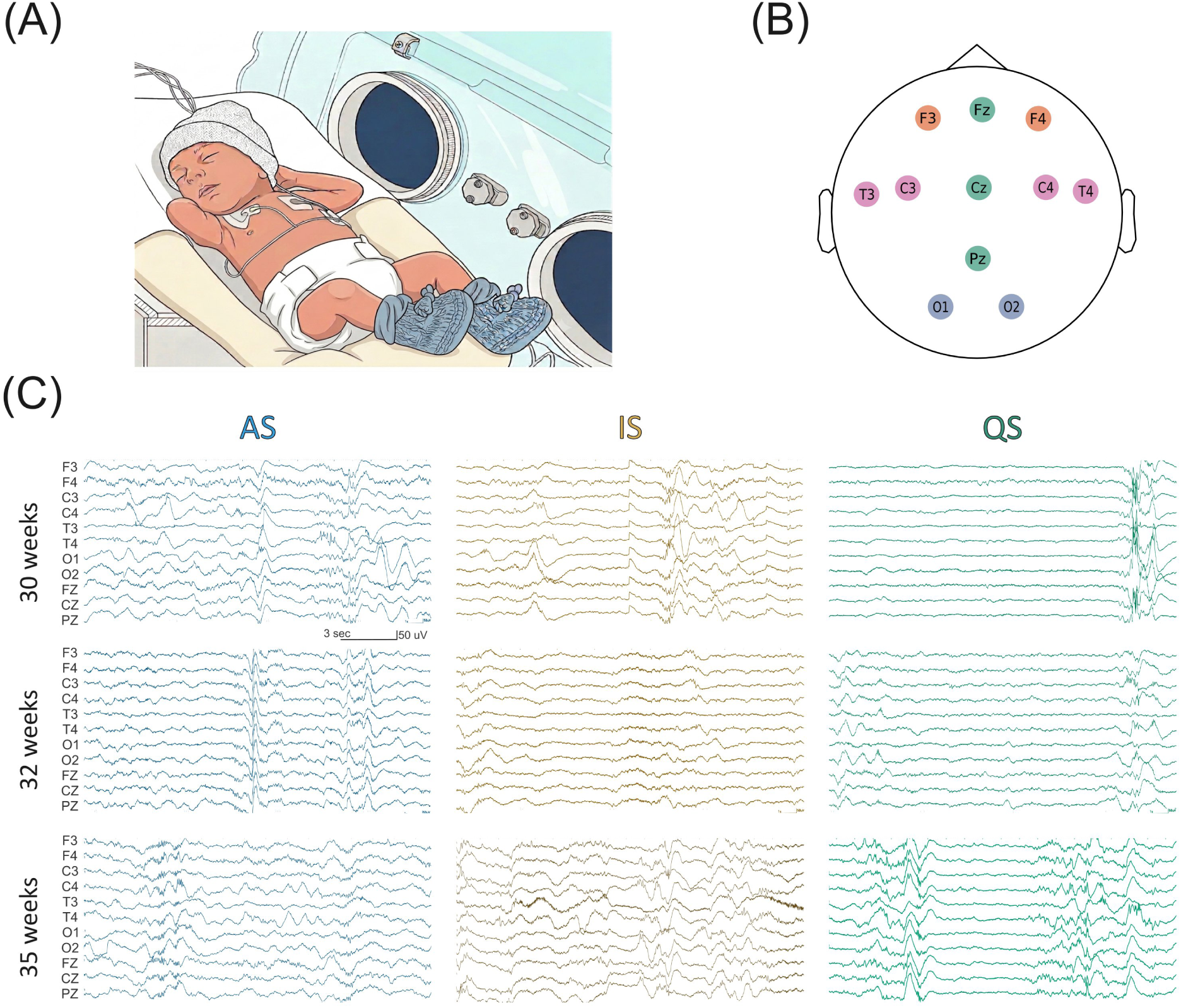
Recording and analysis of preterm infants. A. Representative image of a premature baby during polysomnographic recording. B. Location of the EEG electrodes in the recorded patient. C. Example of EEG recordings at different weeks of postmenstrual age. AS, active sleep; IS, indeterminate sleep; QS, quiet sleep.

### 2.3. Sleep-Wake scoring and analysis

Recordings were acquired using Nihon Kohden’s PortaView system and were subsequently scored manually with PortaView Review software by two trained neurophysiologists. Sleep states were manually scored according to standard criteria based on EEG activity (using longitudinal and transverse bipolar montages), EOG and EMG signals, behavior and vocalizations assessed from synchronized audio–video recordings, and cardiorespiratory parameters (Curzi-Dascalova et al., 1988;Cherian et al., 2009;Andre et al., 2010). Sleep and wake states were classified as AS, QS, IS, or W, in 20-second epochs. AS was characterized by closed eyes, brief facial (smiles, sucking, grimaces, and blinks) and body movements (twitches), isolated vocalizations, intermittent eye movements, variable or augmented heart rate and respiration, continuous EEG activity, and absent or transient mylohyoid muscle tone. QS was defined by eyes closed, scarce body movements, regular heart rate and respiration, presence of mylohyoid tone, and discontinuous EEG pattern. IS was scored when polysomnographic features of both AS and QS were present. W was defined by open eyes, crying or other vocalizations, frequent gross body movements, irregular heart rate and respiration, and continuous EEG activity with movement artifacts.

### 2.4. EEG preprocessing

EEG recordings were converted from the proprietary Nihon Kohden format to European Data Format (EDF) for further analysis in MATLAB (R2015a, MathWorks, Natick, MA, USA) and Python. Thereafter, a 50 Hz notch filter was applied to eliminate power-line noise and the EEG was off-line re-referenced to the mastoids. The recordings were then visually inspected and epochs containing movement artifacts, electrode artifacts, or amplifier saturation were discarded. Each 20-second epoch, which was scored manually, was subdivided into five-second segments that were free of artifacts for complexity analysis. These segments were selected because they provide sufficiently long stationary signals for reliable estimation of complexity, while minimizing the likelihood of residual artefacts. In addition to the raw EEG, low-frequency (1–15.5 Hz) and high-frequency (16–30 Hz) signals were obtained by applying zero-phase Butterworth band-pass filters for subsequent complexity analysis.

### 2.5 EEG non-linear analysis

Due to the marked nonlinearity of EEG signals (Palus, 1996), it is essential to employ mathematical tools capable of capturing these intrinsic nonlinear dynamics. In this context, techniques based on information theory have proven particularly valuable for the analysis of electrophysiological signals (Shannon, 1997). In the present study, we employed two metrics for EEG analysis, LZC and JLZC, with which our group has extensive experience (Pascovich et al., 2022;Gonzalez et al., 2023a;Mondino et al., 2024;Pascovich et al., 2024;Catanzariti et al., 2025).

LZC is an information-theoretic measure inspired by the concept of Kolmogorov complexity, which reflects the minimal amount of information required to describe a sequence, providing an index of signal irregularity and dynamical richness (Lempel and Ziv, 1976). To estimate the complexity of a time series *X*(*t*) ≡ [*x*_t_ ; *t* = 1, ⋯, *T*], we utilized the Lempel and Ziv scheme proposed in 1976. In this approach, a sequence *X*(*t*) is parsed into a number *W* of words by considering any subsequence that has not yet been encountered as a new word. The Lempel-Ziv complexity *c*_LZ_ is the minimum number of words *W* required to reconstruct the information contained in the original time series. For example, the sequence 100110111001010001011 can be parsed in 7 words: 1 · 0 · 01 · 101 · 1100 · 1010 · 001011, giving a complexity *c*_LZ_ = 7. An easy way to apply the Lempel-Ziv algorithm can be found in (Kaspar and Schuster, 1987). The LZC was normalized based on the length *T* of the discrete sequence and the alphabet length (α) as:

LZC = *c*_LZ_ [log_α_*T*] / *T*

Although initially Lempel and Ziv developed the method for binary sequences, the length of the alphabet could be used for any alphabet with finite length. For continuous sequences, such as EEG, the signal must first be discretized; in this work the signals were binarized using the median as threshold.

LZC was computed from both raw EEG signals and signals filtered into lower (1-15.5 Hz) and higher (16-30 Hz) frequency bands, to investigate potential differences in signal complexity across these components (Gonzalez et al., 2022;Catanzariti et al., 2025). The 1-15.5 Hz band encompasses slow and intermediate frequencies that dominate preterm EEG activity and are strongly modulated by maturation and behavioral state. The 16-30 Hz band includes faster oscillatory components that are less prominent in premature infants and are often observed as part of developmental transients such as delta brushes (Vanhatalo and Kaila, 2006;Andre et al., 2010;Kaminska et al., 2018). For each 5-second epoch and frequency band, LZC was computed independently for each EEG channel in every subject. The resulting values were averaged across epochs for each channel, yielding a representative LZC estimate for each channel. Subsequently, grand averages for each channel were computed across subjects for each sleep state. Additionally, the median LZC (mLZC) for all the EEG channels, or for each hemisphere, were also computed and compared across PMA and sleep states.

JLZC. By extending the alphabet size, this approach can be easily generalized to multivariate discrete processes (Zozor et al., 2005). This can be done by considering an *m*-dimensional stationary process **X**(*m*), which generates the sequences *x*_t,i_ = *x*_1,i_, …, *x*_T,i_ with *i* = 1, …, *m*, each one of them from an alphabet of α symbols. In our case, these sequences come from the binarization of the EEG signals.

Let *z*_t_ = *z*_1_, …, *z*_T_ be a new sequence defined over an extended alphabet of size α_m_: *z*t = Σi=1^m^ α^i−1^*x*t,i

Then, the JLZC of the *x*_i_-sequence can be calculated as the complexity of the new sequence *z*_t_: *C*_LZ_(*x*_t,i_) = *C*_LZ_(*z*_t_).

In our case *z*_t_ is a new sequence which represents the information from all recorded channels. The main advantage of the JLZC is that it provides global information about brain dynamics, in contrast to the classical LZC analysis per channel. In this work the JLZC was binarized (α = 2) using the median value.

Intuitively, JLZC provides a proxy measure of the extent to which multiple time series co-vary over time. With this concept in mind, we analyzed JLZC in frontal (F3 and F4), temporal (T3 and T4), and occipital (O1 and O2) cortical regions. In addition, we evaluated JLZC separately for the right (F4, T4, and O2) and left (F3, T3, and O1) hemispheres. JLZC values were then assessed for the different sleep states and PMA groups. The hemispheric grouping enables the assessment of left-right asymmetries in large-scale cortical organization, whereas the frontal, temporal, and occipital configurations probe long-range interactions within each cortical region. By combining these complementary spatial scales, JLZC can be evaluated at different levels of cortical organization, providing a more comprehensive characterization of the spatiotemporal neonatal brain activity during sleep and across maturation. Since this is a relatively novel analytical approach, and its interpretation is less straightforward than that of conventional LZC analyses, it was applied only to the unfiltered (raw) EEG recordings. All LZC and JLZC computations were performed using custom Python scripts implementing the Kaspar and Schuster algorithm (Kaspar and Schuster, 1987).

### 2.6. Statistical Analyses

All recordings were analyzed as independent observations, even though some infants contributed with two or more recordings. Complexity data are presented as the median ± interquartile range (IQR; 25^th^-75^th^ percentile). In addition, selected results are displayed as topoplots to facilitate the visualization and interpretation of the spatial distribution of complexity changes.

Since the distribution of EEG complexity measures did not satisfy normality assumptions and developmental changes were not strictly linear, non-parametric tests were applied. Within-group comparisons were conducted for each EEG channel using the Wilcoxon signed-rank test to assess differences in LZC between AS and QS (IS was not included in these analyses because its values were intermediate between those of AS and QS). In addition, the median LZC across recorded channels (mLZC) was compared among AS, QS, and IS using the Friedman test, followed by Wilcoxon signed-rank *post hoc* tests with Bonferroni correction for multiple comparisons.

We also compared the extreme PMA groups (30-31 vs. 34-35 weeks) for each EEG channel within each sleep state using the Mann-Whitney U test. Furthermore, the mLZC was compared among all PMA groups using the Kruskal–Wallis test, followed by Dunn’s *post hoc* test with Bonferroni correction for multiple comparisons. In addition, Spearman rank correlation analyses were performed to evaluate whether the mLZC exhibited maturational trends across PMA. We also assessed interhemispheric differences in mLZC using the Wilcoxon signed-rank test.

To assess spatial differences in JLZC, a zone-based statistical analysis was conducted. For each vigilance state, JLZC values were compared across cortical zones using the Friedman test, followed by Wilcoxon signed-rank *post hoc* tests with Bonferroni correction for multiple comparisons. Differences between sleep states (AS vs. QS) were assessed using the Wilcoxon signed-rank test.

Statistical significance was set at *p* < 0.05, with Bonferroni correction when appropriate. All statistical analyses were performed in Python using the SciPy library, with *post-hoc* analyses conducted using the scikit *post-hocs* package (Virtanen et al., 2020).

## 3. Results

A total of 54 EEG recordings obtained from 39 preterm infants born between 30 and 35 weeks of PMA were included in the study. Clinical and demographic characteristics are presented in Table 1. At the time of the recordings, none of the infants required mechanical ventilation, 10 were receiving non-invasive ventilatory support as continuous positive airway pressure (CPAP), 33 recordings were under caffeine treatment, and none had clinical or laboratory evidence of ongoing sepsis.

**Table 1.** Demographic and clinical characteristics of the preterm infants.

| **39 infants, 54 recordings** |  |
| --- | --- |
| Gestational age at birth (weeks) | 30 ± 2 |
| Birthweight (g) | 1336 ± 408 |
| Head circumference at birth (cm) | 27.8 ± 2.5 |
| Apgar at 5 min | 8 ± 1 |
| Sex Female/Male (n) | 23/16 |
| Postmenstrual age at recording (weeks) | 33 ± 2 |
| Postnatal age (days) | 23 ± 18 |
| Weight at recording (g) | 1568 ± 388 |
| Maternal age (years) | 28 ± 6 |
| Pregnancy Multiple (n) | 3 |
| Pregnancy Single (n) | 36 |
| Mode of delivery (cesarean/vaginal) (n) | 32/7 |
| Complete antenatal corticosteroids YES/NO (n) | 30/9 |
| CPAP, YES/NO (n) | 33/6 |
| Invasive mechanical ventilatory assistance, YES/NO (n) | 8/31 |
| Caffeine, YES/NO (n) | 23/16 |
| Sepsis, YES/NO (n) | 5/34 |
| Oxygen (days) | 7.6 ± 18.8 |

Continuous variables are presented as mean ± standard deviation; categorical data as “n”. The data for the last five variables refer to the neonatal intensive care unit (NICU) stay rather than the time of EEG recording. The conditions during EEG recording are described in the Results section. CPAP, continuous positive airway pressure.

Figure 1C shows representative EEG recordings during AS, QS, and IS, illustrating the clear maturational changes that occur across PMA. Notably, the duration of QS interburst (quiescent) periods progressively decreased with advancing PMA.

### 3.1 Lempel-Ziv complexity analysis

We applied LZC analysis to the raw (unfiltered) EEG recordings across different sleep states and PMA groups. The results are illustrated in the topoplots shown in Figure 2A. LZC values were consistently higher during QS than during AS across all PMA groups (Figure 2B), whereas IS exhibited intermediate values between those observed in AS and QS. The QS-related increase in LZC was more pronounced over the central region in the most immature infants, while in the 34-35 PMA group it was more evident in the anterior and posterior regions. In addition, the topoplots revealed an increase in LZC with maturation, particularly during AS (Figure 2C).

**Figure 2.**
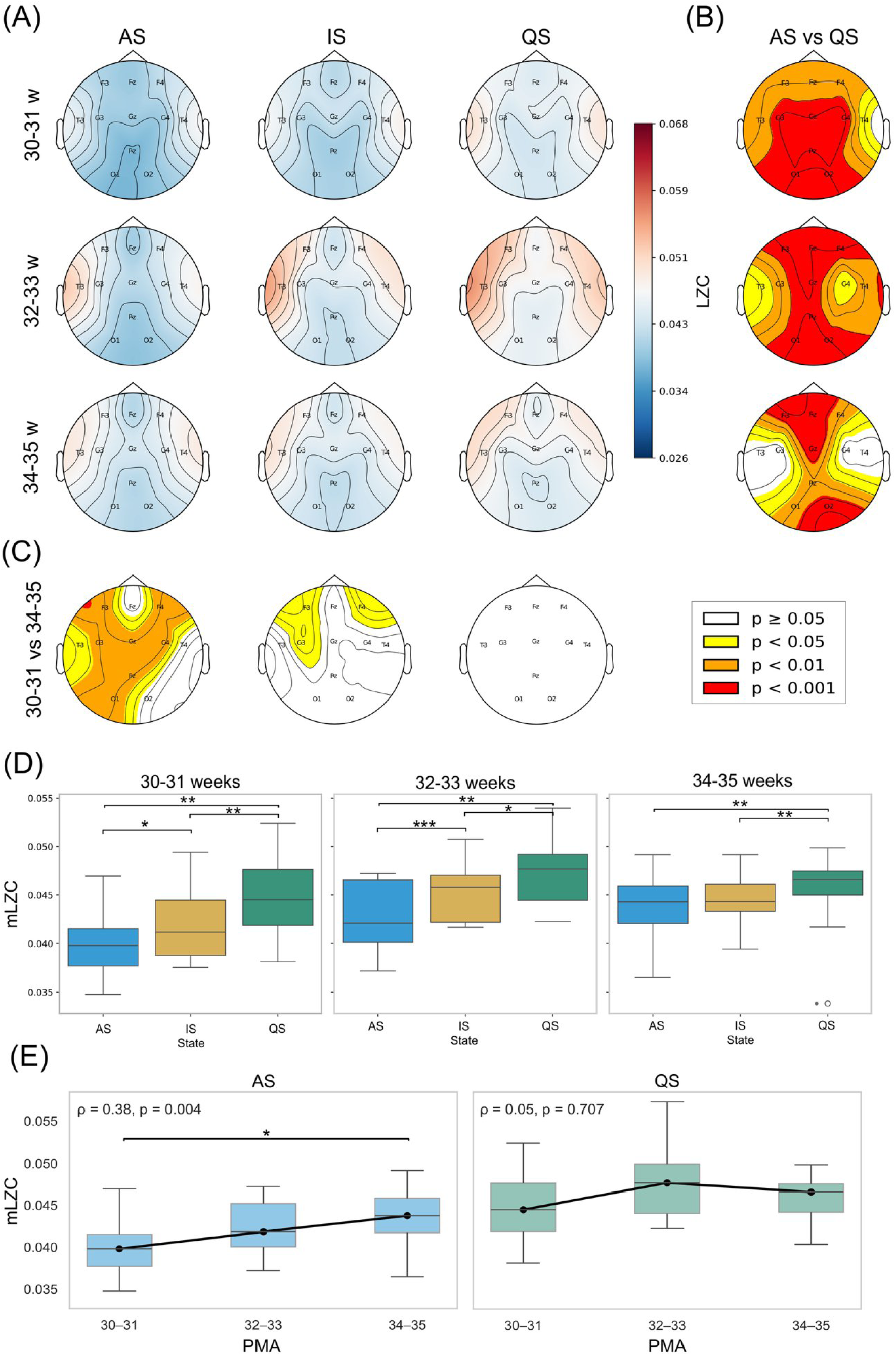
Impact of behavioral states and postmenstrual age on EEG Lempel-Ziv complexity of the raw (unfiltered) recordings. A. Topographic distribution of Lempel-Ziv complexity (LZC) across PMA groups and sleep states. Contour lines represent iso-LZC regions, and electrode locations are indicated. B. Statistical comparison of LZC between active sleep (AS) and quiet sleep (QS) across PMA groups (Wilcoxon signed-rank test). Contour lines denote regions of equal *p*-value. C. Statistical comparison between 30-31 and 34-35 weeks PMA groups within each sleep state (Mann-Whitney U test). D. Boxplots display mLZC across EEG channels for AS, QS, and indeterminate sleep (IS) within each PMA group. Boxes indicate the interquartile range (IQR), the central line represents the median, and whiskers extend to 1.5 × IQR. Statistical comparisons among sleep states were performed using the Friedman and Wilcoxon signed-rank tests. E. Boxplots show the mLZC across EEG channels for the different PMA groups within each sleep state. Spearman correlation coefficients and associated *p*-values are shown to assess maturational trends across PMA. Statistical comparisons among PMA groups were performed using the Kruskal-Wallis and Dunn’s test. In panels D and E: *, *p* < 0.05; **, *p* < 0.01; ***, *p* < 0.001. Number of recordings in each PMA group: 30-31, n = 12; 32-33, n = 16; 34-35 weeks, n = 26.

Interestingly, LZC values in the temporal region were particularly high in the 32-33 weeks PMA group across all sleep states. During AS, the LZC of the temporal-channels in the 32-33 weeks PMA group were significantly higher than that observed in the other PMA groups. For the channel T3, the LZC for the 32-33 weeks group was 0.0500 ± 0.0087; 30-31 weeks 0.0431 ± 0.0069; 34-35 weeks 0.0482 ± 0.0048 (median ± IQR; *p* = 0.03, Kruskal–Wallis and Dunn’s test). Similar results were obtained for T4. During QS, for the channel T3, 32-33 weeks 0.0523 ± 0.0095; 30-31 weeks 0.0471 ± 0.0099; 34-35 weeks 0.0486 ± 0.0056 (*p* = 0.025). Comparable results were obtained for T4.

The mLZC was also significantly higher during QS than during the other sleep states (Figure 2D). Furthermore, mLZC increased with maturation during AS, a finding supported by a significant Spearman rank correlation. In contrast, no significant correlation between mLZC and PMA was observed during QS (Figure 2E).

To obtain a clearer characterization of signal complexity, we analyzed the low (1-15.5 Hz) and high-frequency (16-30 Hz) components of the EEG. The topoplots of Figure 3A represent the LZC of the low-frequency EEG band. The LZC was higher during QS than during AS across several cortical areas in all three PMA groups (Figure 3B). IS exhibited LZC values between those observed during AS and QS. In addition, Figure 3A,C shows that LZC progressively increased with advancing PMA in several cortical areas, particularly during AS.

**Figure 3.**
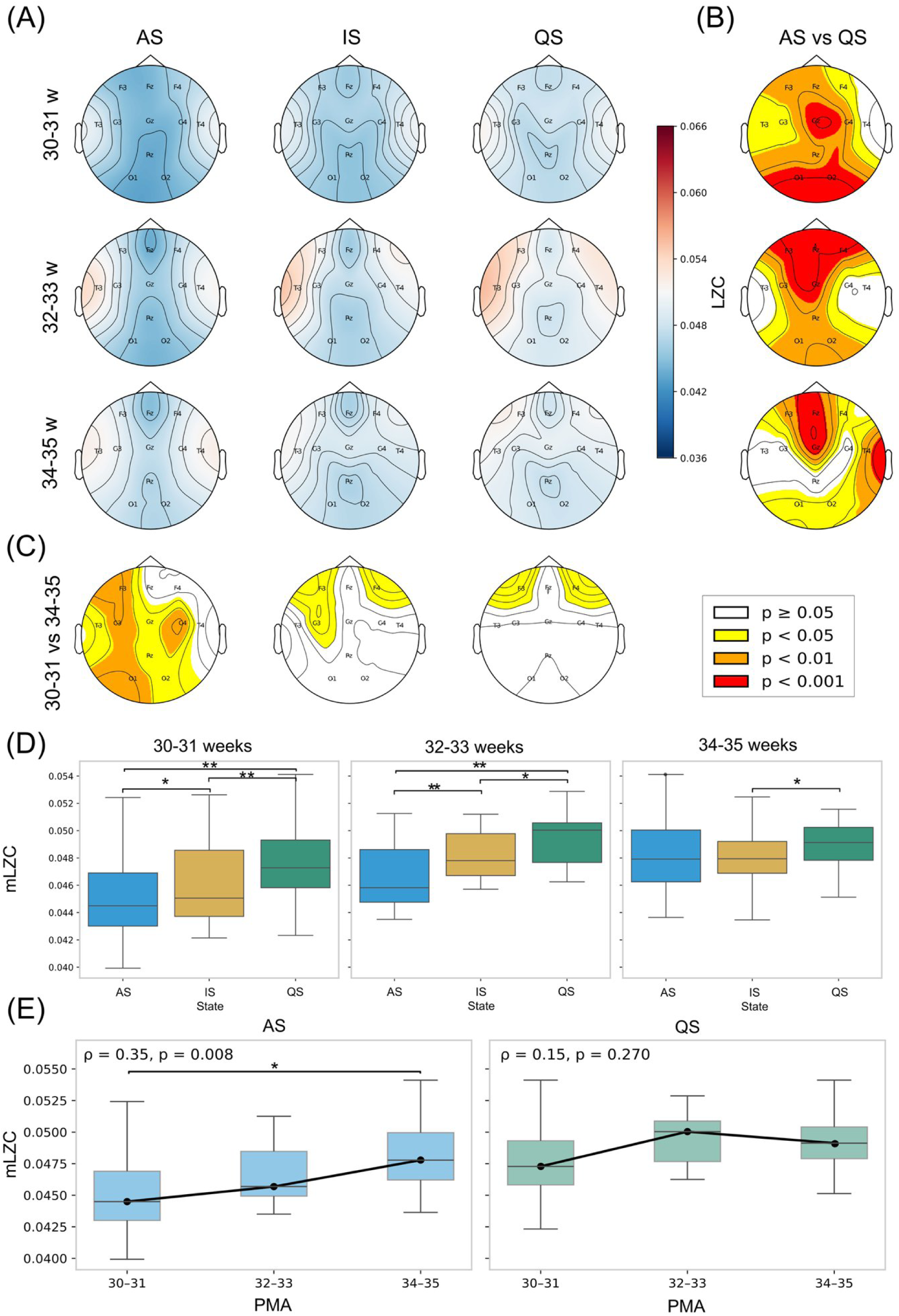
Impact of behavioral states and postmenstrual age on EEG Lempel-Ziv complexity for the low-frequency band (1-15.5 Hz). A. Topographic distribution of Lempel-Ziv complexity (LZC) across PMA groups and sleep states. Contour lines represent iso-LZC regions, and electrode locations are indicated. B. Statistical comparison of LZC between active sleep (AS) and quiet sleep (QS) across PMA groups (Wilcoxon signed-rank test). Contour lines denote regions of equal *p*-value. C. Statistical comparison between 30-31 and 34-35 weeks PMA groups within each sleep state (Mann-Whitney U test). D. Boxplots display mLZC across EEG channels for AS, QS, and indeterminate sleep (IS) within each PMA group. Boxes indicate the interquartile range (IQR), the central line represents the median, and whiskers extend to 1.5 × IQR. Statistical comparisons among sleep states were performed using the Friedman and Wilcoxon signed-rank tests. E. Boxplots show the mLZC across EEG channels for the different PMA groups within each sleep state. Spearman correlation coefficients and associated *p*-values are shown to assess maturational trends across PMA. Statistical comparisons among PMA groups were performed using the Kruskal-Wallis and Dunn’s test. In panels D and E: *, *p* < 0.05; **, *p* < 0.01; ***, *p* < 0.001. Number of recordings in each PMA group: 30-31, n = 12; 32-33, n = 16; 34-35 weeks, n = 26.

As for the raw unfiltered recordings, the LZC values for the low-frequency bands in the temporal region were particularly high in the 32-33 weeks PMA group across all sleep states. During AS, there was a trend in the temporal-channel LZC in the 32-33 weeks PMA to be higher than in the other PMA groups. For channel T3, 32-33 weeks 0.0525 ± 0.0054; 30-31 weeks 0.0481 ± 0.0035; 34-35 weeks 0.0515 ± 0.0041; *p* = 0.06. Similar results were obtained for T4. During QS the LZC in the temporal channel was significantly larger in the 32-33 weeks PMA group; for channel T3, 32-33 weeks 0.0520 ± 0.0043; 30-31 weeks 0.0493 ± 0.0045; 34-35 weeks 0.0501 ± 0.0042; *p* = 0.02. Comparable results were obtained for T4.

The mLZC was also significantly higher during QS than during the other sleep states for the 30-31 and 32-33 PMA groups, while larger than in IS for the 34-35 PMA group (Figure 3D). Furthermore, mLZC increased with maturation during AS, a finding supported by a significant Spearman rank correlation. In contrast, no significant correlation between mLZC and PMA was observed during QS (Figure 3E).

We also performed the same type of analysis for the high frequency band of the EEG. LZC was larger during QS in comparison to AS in most areas in all the PMA groups (Figure 4A and B). The IS has intermediate LZC values. Furthermore, Figure 4A,C shows that in contrast to the LZC of the low-frequency band, LZC of the high-frequency progressively decreased with advancing PMA across several cortical areas in all sleep states; this effect was more evident for AS.

**Figure 4.**
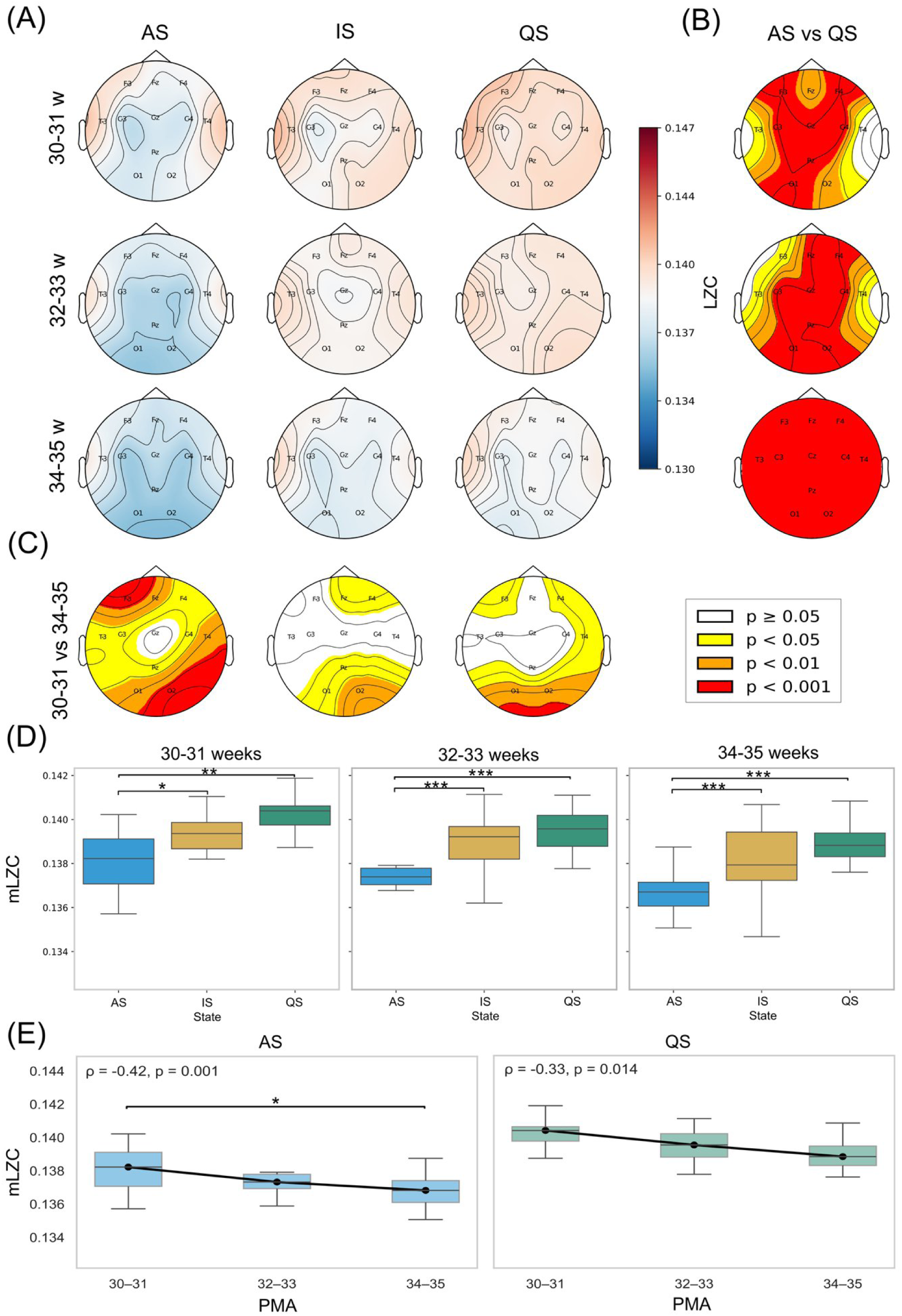
Impact of behavioral states and postmenstrual age on EEG Lempel-Ziv complexity for the high-frequency band (16-30Hz). A. Topographic distribution of Lempel-Ziv complexity (LZC) across PMA groups and sleep states. Contour lines represent iso-LZC regions, and electrode locations are indicated. B. Statistical comparison of LZC between active sleep (AS) and quiet sleep (QS) across PMA groups (Wilcoxon signed-rank test). Contour lines denote regions of equal *p*-value. C. Statistical comparison between 30-31 and 34-35 weeks PMA groups within each sleep state (Mann-Whitney U test). D. Boxplots display mLZC across EEG channels for AS, QS, and indeterminate sleep (IS) within each PMA group. Boxes indicate the interquartile range (IQR), the central line represents the median, and whiskers extend to 1.5 × IQR. Statistical comparisons among sleep states were performed using the Friedman and Wilcoxon signed-rank tests. E. Boxplots show the mLZC across EEG channels for the different PMA groups within each sleep state. Spearman correlation coefficients and associated *p*-values are shown to assess maturational trends across PMA. Statistical comparisons among PMA groups were performed using the Kruskal-Wallis and Dunn’s test. In panels D and E: *, *p* < 0.05; **, *p* < 0.01; ***, *p* < 0.001. Number of recordings in each PMA group: 30-31, n = 12; 32-33, n = 16; 34-35 weeks, n = 26.

The mLZC was also significantly higher during QS than during the other sleep states for all the PMA groups (Figure 4D). Furthermore, mLZC decreased with maturation during AS, a finding supported by a significant Spearman rank correlation. This negative correlation was also observed during QS (Figure 4E).

We also analyzed the mLZC from the raw (unfiltered), low and high-frequency recordings, comparing the right and left hemispheres (Figure 5). Interestingly for the raw and the low-frequency recording the LZC was higher in the left hemisphere for all the sleep states in the 32-33 PMA group. These differences were primarily driven by differences in the temporal channels (T3 and T4), both during AS (*p* = 0.0092) and QS (*p* = 0.00042) in the unfiltered recordings (Wilcoxon signed-rank test). Similar results were obtained for the low-frequency band.

**Figure 5.**
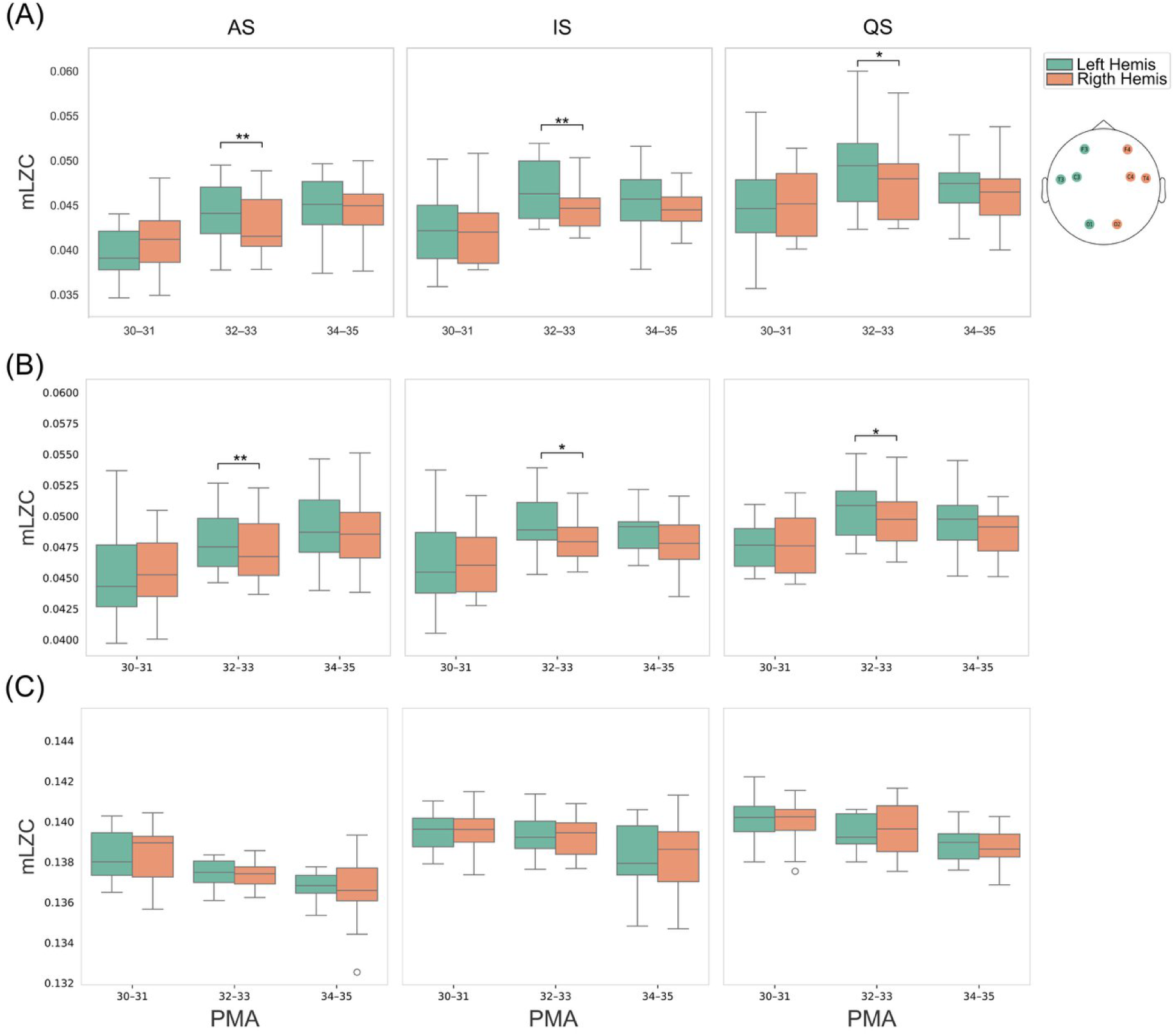
Interhemispheric differences in the Lempel-Ziv complexity. A. Boxplots display mLZC values for the raw (unfiltered) EEG in the right and left hemispheres during active sleep (AS), quiet sleep (QS), and indeterminate sleep (IS) within each PMA group. Boxes indicate the interquartile range (IQR), the central line represents the median, and whiskers extend to 1.5 × IQR. B and C. The same analyses for low and high-frequency EEG channels, respectively. *, *p* < 0.05; **, *p* < 0.01; Wilcoxon signed-rank test . Number of recordings per group: 30-31, n = 12; 32-33, n = 16; 34-35 weeks, n = 26.

### 3.2 Joint Lempel-Ziv complexity analysis

We first compared JLZC across the different cortical regions (Figure 6A). Interestingly, JLZC was consistently highest in the temporal region and lowest in the occipital region across all sleep states and PMA groups. Intermediate values were observed in the frontal region. Similarly, IS exhibited JLZC values between those of AS and QS (data not shown). Comparisons of JLZC across PMA groups for the same sleep state revealed no significant differences in any of the cortical regions analyzed.

**Figure 6.**
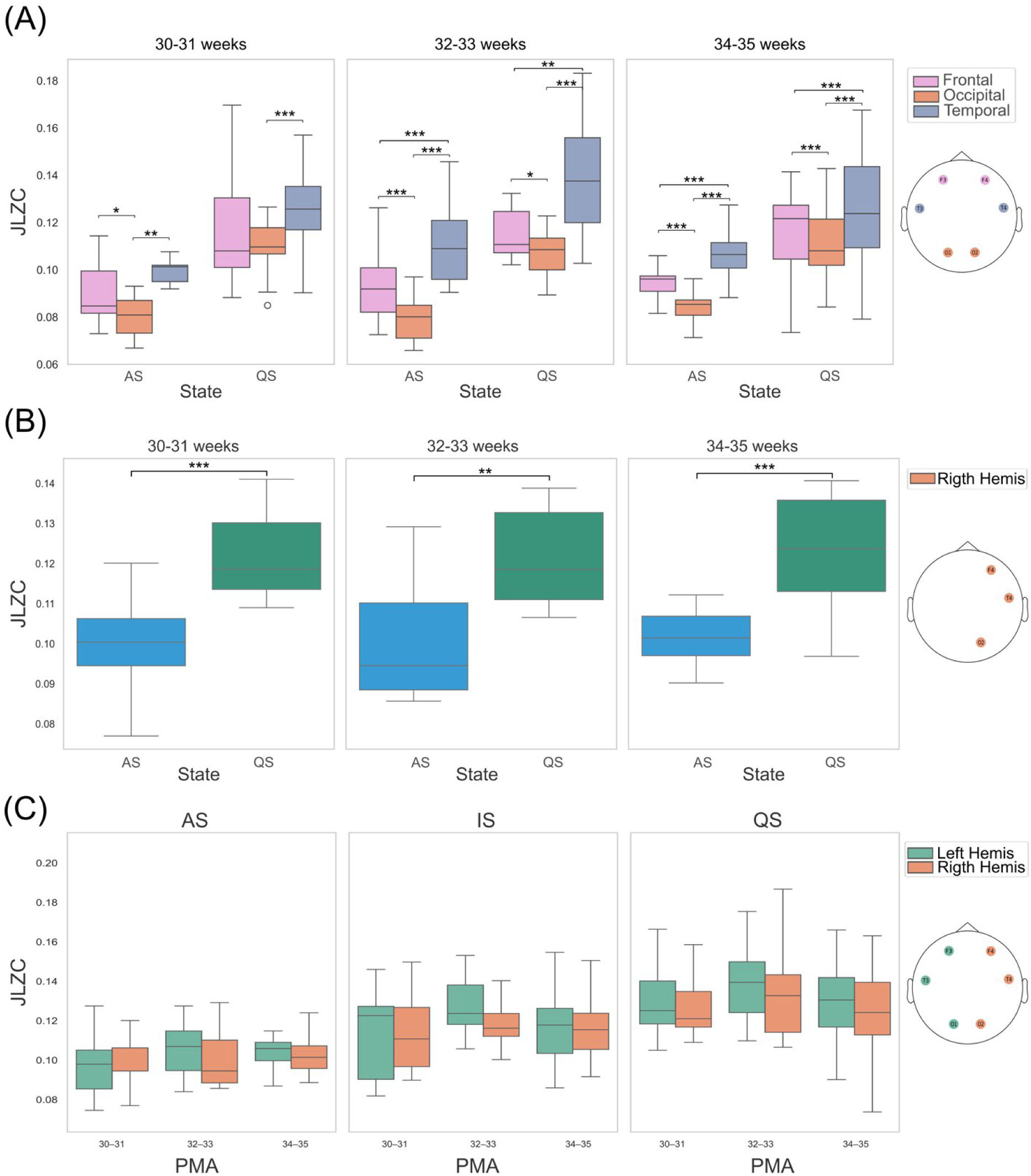
Joint Lempel-Ziv complexity analysis. A. Boxplots display joint Lempel-Ziv complexity (JLZC) for the raw (unfiltered) EEG of in the frontal, temporal, and occipital regions across sleep states and postmenstrual age (PMA) groups. Boxes indicate the interquartile range (IQR), the central line represents the median, and whiskers extend to 1.5 × IQR. Statistical comparisons among sleep states were performed using the Friedman and Wilcoxon signed-rank tests. B. JLZC of the right hemisphere during active sleep (AS) and quiet sleep (QS) across PMA groups. Comparison between sleep states were performed using the Wilcoxon signed-rank test. C. JLZC of the left and right hemispheres across sleep states and PMA groups. Statistical analyses were performed using the Wilcoxon signed-rank test. *, *p* < 0.05; **, *p* < 0.01; ***, *p* < 0.001. Number of recordings per group: 30-31, n = 12; 32-33, n = 16; 34-35 weeks, n = 26. Hemis, hemisphere.

We also evaluated hemispheric JLZC. As shown in Figure 6B, JLZC in the right hemisphere was significantly higher during QS than during AS, whereas IS displayed intermediate values (data not shown). Similar results were observed in the left hemisphere (data not shown). In addition, no significant differences in JLZC were found between the right and left hemispheres across sleep states or PMA groups (Figure 6C). Finally, comparisons of JLZC across PMA groups within the same sleep state revealed no significant differences in either hemisphere.

## 4. Discussion

The functional organization of the developing brain during premature extrauterine life remains a critical yet poorly understood frontier in neonatal neuroscience. In the present study, we evaluated brain dynamical complexity across sleep states in preterm infants using single-channel LZC in different EEG frequency bands. In addition, we applied JLZC, representing, to our knowledge, the first application of this multivariate metric to neonatal EEG recordings. While LZC yielded robust, frequency-dependent markers of cortical maturation and sleep-state differentiation, the introduction of JLZC provides a novel framework to assess the coordinated, multidimensional variability across distinct cortical regions. Together, these non-linear approaches demonstrate that non-invasive electrophysiological complexity metrics can capture both localized microstructural maturation and large-scale functional dynamics during a key developmental window of human brain development.

### 4.1. Methodological considerations

The preterm EEG was analyzed across sleep stages, which become well established from approximately 30 weeks of PMA (Cherian et al., 2009;Andre et al., 2010;Dereymaeker et al., 2017). W was excluded from the analysis, as it accounted for less than 1% of the recordings within the maturational window studied (Devera et al., 2026). In addition, the EEG recordings during W contained numerous artifacts related to limb movements and vocalizations, precluding reliable analysis (Andre et al., 2010).

We excluded preterm patients with severe conditions from the study. However, it is important to consider that preterm infants, even in the absence of obvious brain injury, have an approximately two-fold increased risk of cognitive and behavioral impairments compared with term-born infants (Eeles et al., 2017;Spittle et al., 2017). In fact, brain development differs substantially between healthy fetuses and preterm infants, highlighting the potential influence of the extrauterine environment (Ball et al., 2016;Lefevre et al., 2016). Thus, the concept of “normality” in this population should be interpreted with caution. Moreover, compared with preterm neonates of the same PMA, fetuses spend more time in a sleep-like state, largely due to the influence of sedative substances produced by the placenta and the fetus itself (Mellor et al., 2005;Koch, 2009;Dehaene-Lambertz, 2024). Therefore, although it is challenging to talk about a normal ex utero brain development, it is generally accepted as a close approximation to study normal functional brain maturation (Dereymaeker et al., 2017).

Since this was our first approach to EEG analysis in this population, we did not consider important factors that should be addressed in future studies. Two of these are the duration of exposure to the extrauterine environment (postnatal days) and the gestational age at birth, which on average were 24 ± 19 days and 30 ± 2 weeks, respectively (Table 1). In fact, (Devera et al., 2026) showed that there are subtle changes in sleep architecture according to the postnatal days. A third factor that may have influenced the results is caffeine citrate treatment, as some recordings were acquired while the infants were receiving this medication. Caffeine, a competitive adenosine antagonist that mainly target the A2A receptor, it is used for the treatment of apnea of prematurity and for neuroprotective purposes (Schmidt et al., 2006;O’Shea et al., 2026), but its impact on the ontogeny of brain electrical activity remains unclear. An additional factor is the impact of gender (Frohlich et al., 2024;Smeia et al., 2025); in this regard, the maturation of the GABAergic system, which is critical for the generation of the EEG rhythms, differ between male and female preterm (Lacaille et al., 2021).

### 4.2. Sleep states show different levels of EEG complexity

As a first step toward an in-depth analysis of the preterm EEG, we used the LZC, a metric in which our group has extensive experience (Pascovich et al., 2022;Gonzalez et al., 2023a;Mondino et al., 2024;Pascovich et al., 2024;Catanzariti et al., 2025). First, we analyzed LZC in the raw recordings. To further characterize the results, we analyzed LZC in the low- and high-frequency bands (1-15.5 Hz and 16-30 Hz). The limits of these bands were similar to our previous reports (Gonzalez et al., 2022;Catanzariti et al., 2025), in which, in adult animal models, they correspond to frequency ranges commonly associated with NREM sleep (delta and sigma, including spindle activity) and wakefulness/REM sleep (higher frequencies, particularly the beta band). In preterms, separating EEG activity into low- and high-frequency bands may capture physiologically relevant components of preterm EEG activity, such as the low- and high-frequency components of delta brushes.

The analysis of LZC in the raw EEG signal, as well as in the low- and high-frequency bands, revealed a clear and consistent increase in LZC during QS compared with AS, regardless of PMA. This effect was evident across multiple EEG channels, as illustrated by the topoplots, and was also reflected in the mLZC, a measure of global EEG complexity. LZC values during IS were intermediate between those observed during AS and QS.

These results were, *a priori*, somewhat unexpected, since in adults this metric tends to decrease during adult NREM sleep, as the time series becomes more regular and predictable (Schartner et al., 2017;Pascovich et al., 2022;Mondino et al., 2024;Catanzariti et al., 2025). However, the neurophysiological organization of sleep during prematurity differs substantially from that of the mature brain. In preterm infants, QS is characterized by discontinuous EEG with burst of activity and quiescence periods, prolonged interburst intervals, and highly irregular intra-burst activity rather than the stable synchronized oscillations characteristic of mature NREM sleep (Andre et al., 2010;Grigg-Damberger, 2016). Furthermore, our analysis encompasses the 32-34 weeks PMA period, a developmental window during which transient activity such as delta brushes are fully developed (Wallois et al., 2021). Such temporal discontinuity and heterogeneity may contribute to higher LZC values during QS.

Our findings differ from previous studies in preterm infants, that utilizing different non-linear metrics (multiscale complexity, sample entropy, dimensional complexity, etc.), reported lower values during QS compared with AS (Scher et al., 2005;Janjarasjitt et al., 2008;Zhang et al., 2009;De Wel et al., 2017;De Wel et al., 2019). Several factors may account for these discrepancies. Non-linear metrics may capture distinct properties of neonatal EEG dynamics (Ibáñez-Molina et al., 2026). For example, EEG entropy mainly measures the degree of uncertainty, irregularity, or unpredictability of the signal (Cover, 1999). A signal with high entropy is less predictable, whereas a signal with low entropy is more regular, stereotyped, or synchronized. Therefore, entropy is usually interpreted as a measure of temporal disorder or variability in brain activity. On the other hand, LZC quantifies how much diversity of patterns exists within the signal over time. A signal is considered more complex when many new and non-repetitive patterns appear (Lempel and Ziv, 1976). Thus, complexity is not solely determined by irregularity, but also by the presence of a rich and variable structure, making LZC closely related to sequence diversity and compressibility. Consequently, the discontinuous burst-quiescence activity characteristic of preterm QS may affect these metrics in different ways. Furthermore, it is important to take into account that some studies compare QS with broadly defined non-QS states, a category that includes AS, IS, and likely a small number of W epochs, thereby potentially introducing bias into the results (De Wel et al., 2017). Interestingly, our findings are consistent with those of (Smeia et al., 2025), who reported higher LZC during interburst periods, which are characterized by relative suppression of EEG activity that is more prevalent during QS.

The magnitude and topographic pattern of the differences between QS and AS depended on both PMA and the frequency band analyzed. Nonetheless, it is important to emphasize that, in the high-frequency band, LZC values during QS exceeded those during AS across all EEG channels in the most mature PMA group. One interesting fact is that auditory stimulation elicits delta brushes with frequency components extending into the gamma band (> 31.5 Hz), predominantly during QS (Kaminska et al., 2018). Therefore, exposure to the noisy NICU environment may induce high-frequency activity during this sleep state, potentially contributing to the complexity findings reported here.

### 4.3. Impact of brain maturation on the EEG complexity

In both the raw (unfiltered) EEG signal and the low-frequency band, LZC increased with advancing PMA, but only during AS, which is the predominant sleep state at the studied PMAs (Curzi-Dascalova et al., 1993). Conversely, the high-frequency band exhibited the opposite trend, with LZC progressively decreasing across PMA in both AS and QS.

The LZC increase in the raw recording and low-frequency band was relatively widespread during AS and was also observed in the mLZC. An increase in complexity correlated with PMA has also been reported with LZC or using other complexity measures, when the EEG was analyzed without distinguishing between AS and QS (De Wel et al., 2021;Smeia et al., 2025), during non-QS states (De Wel et al., 2017) or during AS (Scher et al., 2005). In contrast, in the low-frequency band there was a LZC increase during IS and QS localized only in frontal regions, but there was not a significant increase in the mLZC.

As abovementioned the analysis of LZC in the high-frequency band across PMA revealed a progressive decrease with maturation, suggesting that EEG activity becomes more organized and predictable. This reduction in LZC was more spatially widespread during AS, sparing mainly central regions, whereas during both IS and QS the decrease was predominantly observed in frontal and occipital cortical areas. Correlation analyses showed trend consistent with group comparisons.

These changes in LZC across PMA may reflect intrinsic brain maturation and the progressive development of neural network connectivity. In future studies, it would be interesting to investigate whether electrophysiological graphoelements, such as delta brushes, theta waves, and other developmental EEG patterns, whose prevalence and density vary with PMA (Wallois et al., 2021), underlie these findings in EEG complexity.

It is particularly interesting that LZC showed opposite trajectories across PMA in the low- and high-frequency bands. In the adult brain, NREM slow waves depend largely on the intrinsic membrane properties of thalamocortical neurons, which enable them to oscillate in the delta-theta frequency range when hyperpolarized, while the cortex itself is critical to generate slow oscillations (< 1Hz) . In contrast, the beta-gamma activity characteristic of active behavioral states is thought to arise predominantly from cortical circuits, with GABAergic interneurons playing a critical role (Engel and Fries, 2010;Buzsaki and Wang, 2012;Adamantidis et al., 2019;Gonzalez et al., 2023b). Thus, the opposite developmental trajectories observed in low- and high-frequency LZC may reflect the differential maturation of thalamocortical and intracortical mechanisms underlying neonatal sleep states. Interestingly, it has been proposed that the preterm EEG comprises two distinct components with different developmental trajectories: the spontaneous activity transient (SAT) and the apparent continuous oscillatory activity (Vanhatalo and Kaila, 2006). This framework may help explain the different maturational trajectories of LZC that we observed in the low- and high-frequency EEG bands

An important factor in the maturation of EEG rhythms is the developmental evolution of GABAergic neurotransmission. In immature neurons, high intracellular chloride ( *Cl*^−^ *_i_*) levels, sustained by predominant NKCC1 activity relative to low KCC2 cotransporter expression, reverse the transmembrane *Cl*^−^ gradient. As a result, activation of GABA_A_ receptors induces *Cl*^−^ efflux, generating a depolarizing response rather than the hyperpolarizing inhibition typical of the adult brain (Ben-Ari, 2014). This developmental shifts in GABA function and interneuronal network maturation may differentially shape low- and high-frequency EEG dynamics during early brain development.

In addition, non-neuronal factors, such as increased skull thickness and fontanelle closure with maturation, which are known to affect the amplitude and spatial distribution of the brain activity underlying the EEG may affect the LZC (Wallois et al., 2021); however, the impact of these factors remains unknown.

Another issue is the modulatory effects of the extrauterine environment. Notably, LZC values in the temporal region, both in the raw EEG signal and in the low-frequency band (but not in the high-frequency band), were particularly elevated in the 32-33 weeks PMA group across sleep states. Given that the auditory cortex undergoes substantial maturation beginning at approximately 28 weeks of gestation, temporal cortical networks may be especially sensitive to extrauterine auditory experience (Draganova et al., 2007;Toulmin et al., 2015;Adebimpe et al., 2019;Cavalcanti et al., 2020). In fact, both voice and click auditory stimuli evoked cortical responses even before 34 weeks of PMA. At this stage, the responses consisted of a temporal negative slow wave accompanied by rapid oscillations resembling spontaneous delta brushes (Chipaux et al., 2013;Kaminska et al., 2018). Therefore, the increased complexity observed in the temporal region could reflect, at least in part, early experience-dependent functional organization of auditory cortical activity in preterm infants exposed to the acoustic environment of the NICU.

The lateralization of brain functions plays an important role in language processing and attention, and atypical lateralization patterns have been associated with several neurodevelopmental disorders, including autism spectrum disorder (Blanco-Gomez et al., 2025). We detected interhemispheric differences in LZC exclusively in the 32-33 weeks PMA group, with larger LZC values in the left hemisphere for both the raw and low-frequency EEG recordings across sleep states. This transient lateralization, was mainly driven by differences in the temporal cortices, and is consistent with previous reports describing hemispheric asymmetries during this developmental period, including functional connectivity of theta temporal activity coalescing with slow waves (TTA-SW) (Adebimpe et al., 2019), cortical responses to auditory phonemes (Mahmoudzadeh et al., 2013), and the anatomical development of the arcuate fasciculus (Leroy et al., 2015). These findings may reflect early hemispheric specialization that precedes the lateralized brain functions emerging during school age and continuing into adulthood.

### 4.4. Joint Lempel-Ziv complexity

The raw EEG signals were also subjected to JLZC analysis. JLZC provide a metric of coordinated spatiotemporal dynamics and, therefore, reflect certain aspects of functional interactions between cortical regions. In simple terms, this metric provides an estimate of how different signals (EEG channels in this dataset) co-vary over time. For example, if two signals follow a similar temporal pattern, the JLZC between those areas is low; conversely, it is higher when their temporal patterns differ. Overall, temporal regions exhibited higher JLZC values compared with frontal and occipital regions. This pattern was consistently observed across sleep states and PMA. In this regard, these results are consistent with previous reports suggesting that temporal networks undergo reorganization during the period of prematurity. Meijer et al. (2016) demonstrated an age-related decrease in interhemispheric EEG correlation involving temporal leads, indicative of increasing functional differentiation during maturation (Meijer et al., 2016). Interestingly, the resting-state functional connectivity assessed by optical imaging has been shown to be greater between the temporal cortices in preterm infants than in full-term newborns (Fuchino et al., 2013). Although these studies assessed connectivity rather than complexity, they support the notion that temporal cortical regions exhibit distinctive developmental dynamics that may contribute to the elevated JLZC values observed in our cohort.

JLZC in the occipital cortex was lower compared with all other cortical areas. Occipital cortical activity during the preterm period is characterized by the frequent occurrence of SATs, developmental EEG patterns that are thought to contribute to the maturation of thalamocortical and sensory cortical circuits (Wallois et al., 2021). Notably, the frequency of occipital delta brushes increases after approximately 32 weeks PMA, coinciding with a period of rapid visual system development. The repetitive and stereotyped nature of these events may reduce the diversity of joint temporal dynamics, thereby contributing to the lower JLZC values observed in occipital regions. Additionally, methodological factors should also be considered. Because the inter-electrode distance between O1 and O2 is shorter than that between T3 and T4, signal similarity may be increased by volume conduction and overlapping neural sources, potentially contributing to lower JLZC estimates in occipital regions.

We also explored whether developmental differences existed between the two hemispheres. To address this question, we analyzed JLZC separately for the right and left hemispheres. JLZC values were higher during QS than AS, indicating more diverse joint spatiotemporal dynamics across EEG channels during QS. It is important to note that, although QS exhibits stronger long-range and more widespread cortical functional connectivity than AS (Tokariev et al., 2019), and JLZC is influenced by the degree of coordinated temporal dynamics across channels, JLZC does not directly quantify functional connectivity.

JLZC values did not vary significantly across PMA within the same sleep state. These findings differ from those obtained using a measure of functional connectivity, such as coherence or phase lag index, which showed that functional connectivity decreases with increasing PMA (Meijer et al., 2014;van de Pol et al., 2018). Additionally, we found no significant differences in JLZC between the left and right hemispheres for any sleep state or PMA group.

### 4.5. Development of consciousness

Complexity-related measures have been proposed as reliable markers of consciousness across many different conditions, such as sleep, anesthesia, hallucinatory states, coma, and related disorders (Tononi and Edelman, 1998;Sarasso et al., 2021;Frohlich et al., 2022). In fact, in previous studies using animal models, we found reduced complexity during anesthesia (Fuentes et al., 2022;Mondino et al., 2024). LZC also decreased during NREM sleep compared with normal wakefulness; in contrast, REM sleep complexity was higher than that observed during NREM sleep, although still lower than during W (Pascovich et al., 2022;Gonzalez et al., 2023a;Mondino et al., 2024;Catanzariti et al., 2025). In this regard, the study of the emergence of the infant (and fetal) consciousness is a distinctive research focus in the science of consciousness (Bayne et al., 2023). It is considered that the earliest possible time point when subjective experience might begin is likely the establishment of thalamocortical connectivity at 26 weeks of gestation, as the thalamocortical system is necessary for consciousness according to most theoretical frameworks; however, the existence of true consciousness in preterms is still under debate (Padilla and Lagercrantz, 2020;Ciaunica et al., 2021;Bayne et al., 2023;Frohlich et al., 2023;Dehaene-Lambertz, 2024;Delafield-Butt and Ciaunica, 2024;Frohlich and Bayne, 2025). It has been hypothesized that cortical complexity increases as the brain’s capacity for consciousness develops approaching birth, and that this trajectory continues during the neonatal period (Frohlich et al., 2024). Consistent with this hypothesis, we observed an increase in LZC with PMA during AS, the predominant behavioral state at this developmental stage, in both the raw EEG and the low-frequency band. Similar findings have been reported using integrated information analysis of the EEG, a putative measure of consciousness (Isler et al., 2018). In contrast, complexity decreased with PMA in the high-frequency band. A similar decrease in complexity with advancing PMA was reported by Fröhlich et al. (2023b), who analyzed sensory-evoked fetal magnetoencephalography signals using a different analytical approach. Although our study is primarily descriptive, the detailed characterization of LZC and JLZC across sleep states and early developmental stages may provide valuable insights into the neurophysiological processes underlying the emergence of consciousness.

### 4.6. Conclusions and future directions

Understanding the functional maturation of the preterm brain remains a major challenge in neonatal neuroscience. In the present report, we performed a detailed analysis of preterm EEG complexity across PMA and different sleep states in preterm without severe clinical complications at the time of the recordings. The results obtained from LZC, computed in both low- and high-frequency bands, showed a robust effect of sleep state and PMA. In addition, JLZC, a metric that evaluate the degree of coordinated temporal dynamics across areas, revealed that the temporal region, which also exhibited the highest LZC values, displayed significantly different JLZC patterns compared with the frontal and occipital regions. We hypothesize that the elevated values of both complexity measures in the temporal cortex may reflect the influence of extrauterine life during this critical developmental period.

Taken together, these findings provide novel insights into early human neurodevelopment and establish a valuable insight for future clinical and neuroscientific applications. Further analyses using complementary metrics on the same dataset are warranted to extend and refine these observations, including the influence of postnatal age, gestational age at birth, and the potential effects of caffeine treatment.

## Abbreviations

AS: active sleep
CNS: central nervous system
EEG: electroencephalogram
EMG: electromyography
EOG: electrooculography
IQR: interquartile range.
IS: indeterminate sleep
JLZC: joint Lempel-Ziv complexity
LZC: Lempel-Ziv complexity
mLZC: median Lempel-Ziv complexity
NICU: Neonatal Intensive Care Unit
PMA: postmenstrual age
QS: quiet sleep
SAT: spontaneous activity transient.
W: wakefulness

## Acknowledgements

Partially supported by CSIC I+D group 2022 (# 883465) grant and PROINBIO program from Uruguay. In addition, we would like to thank the newborns, their families, and the healthcare staff for making this study possible.

## References

Adamantidis, A.R., Gutierrez Herrera, C., and Gent, T.C. (2019). Oscillating circuitries in the sleeping brain. Nat Rev Neurosci 20, 746–762.

Adebimpe, A., Routier, L., and Wallois, F. (2019). Preterm Modulation of Connectivity by Endogenous Generators: The Theta Temporal Activities in Coalescence with Slow Waves. Brain Topogr 32, 762–772.

Andre, M., Lamblin, M.D., D’allest, A.M., Curzi-Dascalova, L., Moussalli-Salefranque, F., T, S.N.T., Vecchierini-Blineau, M.F., Wallois, F., Walls-Esquivel, E., and Plouin, P. (2010). Electroencephalography in premature and full-term infants. Developmental features and glossary. Neurophysiol Clin 40, 59–124.

Ball, G., Aljabar, P., Arichi, T., Tusor, N., Cox, D., Merchant, N., Nongena, P., Hajnal, J.V., Edwards, A.D., and Counsell, S.J. (2016). Machine-learning to characterise neonatal functional connectivity in the preterm brain. Neuroimage 124, 267–275.

Bayne, T., Frohlich, J., Cusack, R., Moser, J., and Naci, L. (2023). Consciousness in the cradle: on the emergence of infant experience. Trends Cogn Sci 27, 1135–1149.

Ben-Ari, Y. (2014). The GABA excitatory/inhibitory developmental sequence: a personal journey. Neuroscience 279, 187–219.

Bertelle, V., Mabin, D., Adrien, J., and Sizun, J. (2005). Sleep of preterm neonates under developmental care or regular environmental conditions. Early Hum Dev 81, 595–600.

Blanco-Gomez, G., O’reilly, C., Webb, S.J., Elsabbagh, M., and Team, B. (2025). The Development of Lateralized Brain Oscillations in Infants: Lessons From Autism. Dev Psychobiol 67, e70101.

Blumberg, M.S., Dooley, J.C., and Tiriac, A. (2022). Sleep, plasticity, and sensory neurodevelopment. Neuron 110, 3230–3242.

Burlando, G., Uccella, S., Marazzotta, V., Wang, S.H., Palva, J.M., Roascio, M., Rossi, A., Ramenghi, L.A., Nobili, L., and Arnulfo, G. (2026). Quantifying Cortical Maturational Aspects During Different Vigilance States in Preterm Infants by Advanced EEG Analysis. *J Sleep Res*, e70308.

Buzsaki, G., and Wang, X.J. (2012). Mechanisms of gamma oscillations. Annu Rev Neurosci 35, 203–225.

Catanzariti, M., Mondino, A., Torterolo, P., Aimar, H., Olby, N.J., and Mateos, D.M. (2025). Study of changes in brain dynamics during sleep cycles in dogs under effect of trazodone. PLoS One 20, e0335159.

Cavalcanti, H.G., Da Silva Nunes, A.D., Da Cunha, B.K.S., De Freitas Alvarenga, K., Balen, S.A., and Pereira, A., Jr. (2020). Early exposure to environment sounds and the development of cortical auditory evoked potentials of preterm infants during the first 3 months of life. BMC Res Notes 13, 303.

Cherian, P.J., Swarte, R.M., and Visser, G.H. (2009). Technical standards for recording and interpretation of neonatal electroencephalogram in clinical practice. Ann Indian Acad Neurol 12, 58–70.

Chipaux, M., Colonnese, M.T., Mauguen, A., Fellous, L., Mokhtari, M., Lezcano, O., Milh, M., Dulac, O., Chiron, C., Khazipov, R., and Kaminska, A. (2013). Auditory stimuli mimicking ambient sounds drive temporal “delta-brushes” in premature infants. PLoS One 8, e79028.

Ciaunica, A., Safron, A., and Delafield-Butt, J. (2021). Back to square one: the bodily roots of conscious experiences in early life. Neurosci Conscious 2021, niab037.

Cover, T.M. (1999). Elements of information theory. John Wiley & Sons.

Curzi-Dascalova, L., Figueroa, J.M., Eiselt, M., Christova, E., Virassamy, A., D’allest, A.M., Guimaraes, H., Gaultier, C., and Dehan, M. (1993). Sleep state organization in premature infants of less than 35 weeks’ gestational age. Pediatr Res 34, 624–628.

Curzi-Dascalova, L., Peirano, P., and Morel-Kahn, F. (1988). Development of sleep states in normal premature and full-term newborns. Dev Psychobiol 21, 431–444.

De Groot, E.R., Dudink, J., and Austin, T. (2024). Sleep as a driver of pre- and postnatal brain development. Pediatr Res 96, 1503–1509.

De Wel, O., Lavanga, M., Caicedo, A., Jansen, K., Naulaers, G., and Van Huffel, S. (2019). Decomposition of a Multiscale Entropy Tensor for Sleep Stage Identification in Preterm Infants. Entropy (Basel) 21.

De Wel, O., Lavanga, M., Dorado, A.C., Jansen, K., Dereymaeker, A., Naulaers, G., and Van Huffel, S. (2017). Complexity Analysis of Neonatal EEG Using Multiscale Entropy: Applications in Brain Maturation and Sleep Stage Classification. Entropy (Basel) 19.

De Wel, O., Van Huffel, S., Lavanga, M., Jansen, K., Dereymaeker, A., Dudink, J., Gui, L., Huppi, P.S., De Vries, L.S., Naulaers, G., Benders, M., and Tataranno, M.L. (2021). Relationship Between Early Functional and Structural Brain Developments and Brain Injury in Preterm Infants. Cerebellum 20, 556–568.

Dehaene-Lambertz, G. (2024). Perceptual Awareness in Human Infants: What is the Evidence? J Cogn Neurosci 36, 1599–1609.

Delafield-Butt, J., and Ciaunica, A. (2024). Sensorimotor foundations of self-consciousness in utero. Current Opinion in Behavioral Sciences 59, 101428.

Dereymaeker, A., Pillay, K., Vervisch, J., De Vos, M., Van Huffel, S., Jansen, K., and Naulaers, G. (2017). Review of sleep-EEG in preterm and term neonates. Early Hum Dev 113, 87–103.

Devera, A., Legnani, M., Mesquita, C., Urban, L., Gonzalez, J., Hackembruch, J., Mateos, D., Blasina, F., and Torterolo, P. (2026). Sleep-Wake Dynamics in Preterm Infants. Impact of gestational age, gestational age at birth and postnatal days. In preparation.

Devera, A., Legnani, M., Mezquita, C., Urban, L., Gonzalez, J., Hackembruch, J., Blasina, F., and Torterolo, P. (2023). Sleep-wake patterns in preterm newborns: Effect of extrauterine development. IBRO Neuroscience Reports 15, S733.

Draganova, R., Eswaran, H., Murphy, P., Lowery, C., and Preissl, H. (2007). Serial magnetoencephalographic study of fetal and newborn auditory discriminative evoked responses. Early Hum Dev 83, 199–207.

Eeles, A.L., Walsh, J.M., Olsen, J.E., Cuzzilla, R., Thompson, D.K., Anderson, P.J., Doyle, L.W., Cheong, J.L.Y., and Spittle, A.J. (2017). Continuum of neurobehaviour and its associations with brain MRI in infants born preterm. BMJ Paediatr Open 1, e000136.

Engel, A.K., and Fries, P. (2010). Beta-band oscillations--signalling the status quo? Curr Opin Neurobiol 20, 156–165.

Frank, M.G. (2017). Sleep and plasticity in the visual cortex: more than meets the eye. Curr Opin Neurobiol 44, 8–12.

Frohlich, J., and Bayne, T. (2025). Markers of consciousness in infants: Towards a ’cluster-based’ approach. Acta Paediatr 114, 285–291.

Frohlich, J., Bayne, T., Crone, J.S., Dallavecchia, A., Kirkeby-Hinrup, A., Mediano, P.a.M., Moser, J., Talar, K., Gharabaghi, A., and Preissl, H. (2023). Not with a “zap” but with a “beep”: Measuring the origins of perinatal experience. Neuroimage 273, 120057.

Frohlich, J., Chiang, J.N., Mediano, P.a.M., Nespeca, M., Saravanapandian, V., Toker, D., Dell’italia, J., Hipp, J.F., Jeste, S.S., Chu, C.J., Bird, L.M., and Monti, M.M. (2022). Neural complexity is a common denominator of human consciousness across diverse regimes of cortical dynamics. Commun Biol 5, 1374.

Frohlich, J., Moser, J., Sippel, K., Mediano, P.A., Preissl, H., and Gharabaghi, A. (2024). Sex differences in prenatal development of neural complexity in the human brain. Nat. Mental Health 2, 401–416.

Fuchino, Y., Naoi, N., Shibata, M., Niwa, F., Kawai, M., Konishi, Y., Okanoya, K., and Myowa-Yamakoshi, M. (2013). Effects of preterm birth on intrinsic fluctuations in neonatal cerebral activity examined using optical imaging. PLoS One 8, e67432.

Fuentes, N., Garcia, A., Guevara, R., Orofino, R., and Mateos, D.M. (2022). Complexity of Brain Dynamics as a Correlate of Consciousness in Anaesthetized Monkeys. Neuroinformatics 20, 1041–1054.

Gonzalez, J., Cavelli, M., Mondino, A., Pascovich, C., Castro-Zaballa, S., Torterolo, P., and Rubido, N. (2019). Decreased electrocortical temporal complexity distinguishes sleep from wakefulness. Sci Rep 9, 18457.

Gonzalez, J., Cavelli, M., Tort, A.B.L., Torterolo, P., and Rubido, N. (2023a). Sleep disrupts complex spiking dynamics in the neocortex and hippocampus. PLoS One 18, e0290146.

Gonzalez, J., Mateos, D., Cavelli, M., Mondino, A., Pascovich, C., Torterolo, P., and Rubido, N. (2022). Low frequency oscillations drive EEG’s complexity changes during wakefulness and sleep. Neuroscience 494, 1–11.

Gonzalez, J., Torterolo, P., and Tort, A.B.L. (2023b). Mechanisms and functions of respiration-driven gamma oscillations in the primary olfactory cortex. Elife 12.

Grigg-Damberger, M.M. (2016). The Visual Scoring of Sleep in Infants 0 to 2 Months of Age. J Clin Sleep Med 12, 429–445.

Hobson, J.A. (2009). REM sleep and dreaming: towards a theory of protoconsciousness. Nat Rev Neurosci 10, 803–813.

Ibáñez-Molina, A.J., Iglesias-Parro, S., Gálvez-Garzón, M.C., and Soriano, M.F. (2026). Cognitive Processing and EEG Complexity. Entropy (Basel) 28, 761.

Isler, J.R., Stark, R.I., Grieve, P.G., Welch, M.G., and Myers, M.M. (2018). Integrated information in the EEG of preterm infants increases with family nurture intervention, age, and conscious state. PLoS One 13, e0206237.

Janjarasjitt, S., Scher, M.S., and Loparo, K.A. (2008). Nonlinear dynamical analysis of the neonatal EEG time series: the relationship between sleep state and complexity. Clin Neurophysiol 119, 1812–1823.

Kaminska, A., Delattre, V., Laschet, J., Dubois, J., Labidurie, M., Duval, A., Manresa, A., Magny, J.F., Hovhannisyan, S., Mokhtari, M., Ouss, L., Boissel, A., Hertz-Pannier, L., Sintsov, M., Minlebaev, M., Khazipov, R., and Chiron, C. (2018). Cortical Auditory-Evoked Responses in Preterm Neonates: Revisited by Spectral and Temporal Analyses. Cereb Cortex 28, 3429–3444.

Kaspar, F., and Schuster, H.G. (1987). Easily calculable measure for the complexity of spatiotemporal patterns. Phys Rev A Gen Phys 36, 842–848.

Koch, C. (2009). When Does Consciousness Arise? Scientific american 20, 20–21.

Kostovic, I., Sedmak, G., and Judas, M. (2019). Neural histology and neurogenesis of the human fetal and infant brain. Neuroimage 188, 743–773.

Kratimenos, P., Sanidas, G., Simonti, G., Byrd, C., and Gallo, V. (2025). The shifting landscape of the preterm brain. Neuron 113, 2042–2064.

Lacaille, H., Vacher, C.M., and Penn, A.A. (2021). Preterm Birth Alters the Maturation of the GABAergic System in the Human Prefrontal Cortex. Front Mol Neurosci 14, 827370.

Lefevre, J., Germanaud, D., Dubois, J., Rousseau, F., De Macedo Santos, I., Angleys, H., Mangin, J.F., Huppi, P.S., Girard, N., and De Guio, F. (2016). Are Developmental Trajectories of Cortical Folding Comparable Between Cross-sectional Datasets of Fetuses and Preterm Newborns? Cereb Cortex 26, 3023–3035.

Lempel, A., and Ziv, J. (1976). On the complexity of finite sequences. IEEE Transactions on Information Theory 22, 75–81.

Leroy, F., Cai, Q., Bogart, S.L., Dubois, J., Coulon, O., Monzalvo, K., Fischer, C., Glasel, H., Van Der Haegen, L., Benezit, A., Lin, C.P., Kennedy, D.N., Ihara, A.S., Hertz-Pannier, L., Moutard, M.L., Poupon, C., Brysbaert, M., Roberts, N., Hopkins, W.D., Mangin, J.F., and Dehaene-Lambertz, G. (2015). New human-specific brain landmark: the depth asymmetry of superior temporal sulcus. Proc Natl Acad Sci U S A 112, 1208–1213.

Lopes Da Silva, F. (2010). “EEG: origin and measurement,” in EEG-fMRI, eds. C. Mulert & L. Lemieuz. Springer-Verlag), 19–38.

Mahmoudzadeh, M., Dehaene-Lambertz, G., Fournier, M., Kongolo, G., Goudjil, S., Dubois, J., Grebe, R., and Wallois, F. (2013). Syllabic discrimination in premature human infants prior to complete formation of cortical layers. Proc Natl Acad Sci U S A 110, 4846–4851.

Mateos, D.M., Guevara Erra, R., Wennberg, R., and Perez Velazquez, J.L. (2018). Measures of entropy and complexity in altered states of consciousness. Cogn Neurodyn 12, 73–84.

Mateos, D.M., Wennberg, R., Guevara, R., and Perez Velazquez, J.L. (2017). Consciousness as a global property of brain dynamic activity. Phys Rev E 96, 062410.

Meijer, E.J., Hermans, K.H., Zwanenburg, A., Jennekens, W., Niemarkt, H.J., Cluitmans, P.J., Van Pul, C., Wijn, P.F., and Andriessen, P. (2014). Functional connectivity in preterm infants derived from EEG coherence analysis. Eur J Paediatr Neurol 18, 780–789.

Meijer, E.J., Niemarkt, H.J., Raaijmakers, I.P., Mulder, A.M., Van Pul, C., Wijn, P.F., and Andriessen, P. (2016). Interhemispheric connectivity estimated from EEG time-correlation analysis in preterm infants with normal follow-up at age of five. Physiol Meas 37, 2286–2298.

Mellor, D.J., Diesch, T.J., Gunn, A.J., and Bennet, L. (2005). The importance of ’awareness’ for understanding fetal pain. Brain Res Brain Res Rev 49, 455–471.

Mirmiran, M. (1995). The function of fetal/neonatal rapid eye movement sleep. Behav Brain Res 69, 13–22.

Mirmiran, M., Maas, Y.G., and Ariagno, R.L. (2003). Development of fetal and neonatal sleep and circadian rhythms. Sleep Med Rev 7, 321–334.

Mondino, A., Gonzalez, J., Li, D., Mateos, D., Osorio, L., Cavelli, M., Castro-Nin, J.P., Serantes, D., Costa, A., Vanini, G., Mashour, G.A., and Torterolo, P. (2024). Urethane anaesthesia exhibits neurophysiological correlates of unconsciousness and is distinct from sleep. Eur J Neurosci 59, 483–501.

Moser, J., Bensaid, S., Kroupi, E., Schleger, F., Wendling, F., Ruffini, G., and Preissl, H. (2019). Evaluating Complexity of Fetal MEG Signals: A Comparison of Different Metrics and Their Applicability. Front Syst Neurosci 13, 23.

O’shea, M., Butler, L., Holohan, S., Healy, K., O’farrell, R., Shamit, A., Cusack, R., Elhadi, M., Lynch, S., Gilcrest, M., Semberova, J., Branagan, A., O’dea, M.I., Duddy, P., Ambalavanan, N., Allegaert, K., Bearer, C.F., Meehan, J., and Molloy, E.J. (2026). Caffeine and preterm infants: multiorgan effects and therapeutic creep: scope to optimise dose and timing. Pediatr Res 99, 462–469.

Padilla, N., and Lagercrantz, H. (2020). Making of the mind. Acta Paediatr 109, 883–892.

Palus, M. (1996). Nonlinearity in normal human EEG: cycles, temporal asymmetry, nonstationarity and randomness, not chaos. Biol Cybern 75, 389–396.

Park, J. (2020). Sleep Promotion for Preterm Infants in the NICU. Nurs Womens Health 24, 24–35.

Parmelee, A.H., Jr., Wenner, W.H., Akiyama, Y., Schultz, M., and Stern, E. (1967). Sleep states in premature infants. Dev Med Child Neurol 9, 70–77.

Pascovich, C., Castro-Zaballa, S., Mediano, P.a.M., Bor, D., Canales-Johnson, A., Torterolo, P., and Bekinschtein, T.A. (2022). Ketamine and sleep modulate neural complexity dynamics in cats. Eur J Neurosci 55, 1584–1600.

Pascovich, C., Serantes, D., Rodriguez, A., Mateos, D., Gonzalez, J., Gallo, D., Rivas, M., Devera, A., Lagos, P., Rubido, N., and Torterolo, P. (2024). Dorsal and median raphe neuronal firing dynamics characterized by nonlinear measures. PLoS Comput Biol 20, e1012111.

Peirano, P.D., and Algarin, C.R. (2007). Sleep in brain development. Biol Res 40, 471–478.

Riggins, T., Ratliff, E.L., Horger, M.N., and Spencer, R.M.C. (2024). The importance of sleep for the developing brain. Curr Sleep Med Rep 10, 437–446.

Sarasso, S., Casali, A.G., Casarotto, S., Rosanova, M., Sinigaglia, C., and Massimini, M. (2021). Consciousness and complexity: a consilience of evidence. Neurosci Conscious 2021, niab023.

Schartner, M., Seth, A., Noirhomme, Q., Boly, M., Bruno, M.A., Laureys, S., and Barrett, A. (2015). Complexity of Multi-Dimensional Spontaneous EEG Decreases during Propofol Induced General Anaesthesia. PLoS One 10, e0133532.

Schartner, M.M., Pigorini, A., Gibbs, S.A., Arnulfo, G., Sarasso, S., Barnett, L., Nobili, L., Massimini, M., Seth, A.K., and Barrett, A.B. (2017). Global and local complexity of intracranial EEG decreases during NREM sleep. Neurosci Conscious 2017, niw022.

Scher, M.S. (2008). Ontogeny of EEG-sleep from neonatal through infancy periods. Sleep Med 9, 615–636.

Scher, M.S., Waisanen, H., Loparo, K., and Johnson, M.W. (2005). Prediction of neonatal state and maturational change using dimensional analysis. J Clin Neurophysiol 22, 159–165.

Schmidt, B., Roberts, R.S., Davis, P., Doyle, L.W., Barrington, K.J., Ohlsson, A., Solimano, A., Tin, W., and Caffeine for Apnea of Prematurity Trial, G. (2006). Caffeine therapy for apnea of prematurity. N Engl J Med 354, 2112–2121.

Shannon, C.E. (1997). The mathematical theory of communication. 1963. MD Comput 14, 306–317.

Smeia, L., Zaylaa, A., Metaxas, D., Nourhashemi, M., Mahmoudzadeh, M., Birkenfeld, A.L., Sippel, K., Mediano, P.a.M., Preissl, H., Wallois, F., and Frohlich, J. (2025). Neural complexity in preterm infants is predicted by developmental variables. Plos Complex Systems 2, e0000056.

Spittle, A.J., Walsh, J.M., Potter, C., Mcinnes, E., Olsen, J.E., Lee, K.J., Anderson, P.J., Doyle, L.W., and Cheong, J.L. (2017). Neurobehaviour at term-equivalent age and neurodevelopmental outcomes at 2 years in infants born moderate-to-late preterm. Dev Med Child Neurol 59, 207–215.

Tokariev, A., Roberts, J.A., Zalesky, A., Zhao, X., Vanhatalo, S., Breakspear, M., and Cocchi, L. (2019). Large-scale brain modes reorganize between infant sleep states and carry prognostic information for preterms. Nat Commun 10, 2619.

Tononi, G., and Edelman, G.M. (1998). Consciousness and complexity. Science 282, 1846–1851.

Torterolo, P., Castro-Zaballa, S., Cavelli, M., and Gonzalez, J. (2019). “Arousal and normal conscious cognition,” in Arousal in Neurological and Psychiatric Diseases, ed. E. Garcia-Rill. Elsevier), 1–24.

Toulmin, H., Beckmann, C.F., O’muircheartaigh, J., Ball, G., Nongena, P., Makropoulos, A., Ederies, A., Counsell, S.J., Kennea, N., Arichi, T., Tusor, N., Rutherford, M.A., Azzopardi, D., Gonzalez-Cinca, N., Hajnal, J.V., and Edwards, A.D. (2015). Specialization and integration of functional thalamocortical connectivity in the human infant. Proc Natl Acad Sci U S A 112, 6485–6490.

Tsuchida, T.N., Wusthoff, C.J., Shellhaas, R.A., Abend, N.S., Hahn, C.D., Sullivan, J.E., Nguyen, S., Weinstein, S., Scher, M.S., Riviello, J.J., Clancy, R.R., and American Clinical Neurophysiology Society Critical Care Monitoring, C. (2013). American clinical neurophysiology society standardized EEG terminology and categorization for the description of continuous EEG monitoring in neonates: report of the American Clinical Neurophysiology Society critical care monitoring committee. J Clin Neurophysiol 30, 161–173.

Van De Pol, L.A., Van ’T Westende, C., Zonnenberg, I., Koedam, E., Van Rossum, I., De Haan, W., Steenweg, M., Van Straaten, E.C., and Stam, C.J. (2018). Strong Relation Between an EEG Functional Connectivity Measure and Postmenstrual Age: A New Potential Tool for Measuring Neonatal Brain Maturation. Front Hum Neurosci 12, 286.

Vanhatalo, S., and Kaila, K. (2006). Development of neonatal EEG activity: from phenomenology to physiology. Semin Fetal Neonatal Med 11, 471–478.

Virtanen, P., Gommers, R., Oliphant, T.E., Haberland, M., Reddy, T., Cournapeau, D., Burovski, E., Peterson, P., Weckesser, W., Bright, J., Van Der Walt, S.J., Brett, M., Wilson, J., Millman, K.J., Mayorov, N., Nelson, A.R.J., Jones, E., Kern, R., Larson, E., Carey, C.J., Polat, I., Feng, Y., Moore, E.W., Vanderplas, J., Laxalde, D., Perktold, J., Cimrman, R., Henriksen, I., Quintero, E.A., Harris, C.R., Archibald, A.M., Ribeiro, A.H., Pedregosa, F., Van Mulbregt, P., and Scipy, C. (2020). SciPy 1.0: fundamental algorithms for scientific computing in Python. Nat Methods 17, 261–272.

Wallois, F., Routier, L., Heberle, C., Mahmoudzadeh, M., Bourel-Ponchel, E., and Moghimi, S. (2021). Back to basics: the neuronal substrates and mechanisms that underlie the electroencephalogram in premature neonates. Neurophysiol Clin 51, 5–33.

World-Health-Organization (2023). Preterm birth [Online]. Available: https://www.who.int/news-room/fact-sheets/detail/preterm-birth [Accessed].

Zhang, D., Ding, H., Liu, Y., Zhou, C., Ding, H., and Ye, D. (2009). Neurodevelopment in newborns: a sample entropy analysis of electroencephalogram. Physiol Meas 30, 491–504.

Zozor, S., Ravier, P., and Buttelli, O. (2005). On Lempel–Ziv complexity for multidimensional data analysis. Physica A: Statistical Mechanics and its Applications 345, 258–302.

